# Root hairs and mycorrhiza represent alternative phylogenetically conserved strategies for belowground absorptive surface maximization

**DOI:** 10.64898/2026.05.13.723781

**Authors:** Joana Bergmann, Tom Lachaise, Karla M. Barfuss, Emma Bretherick, Elsa Matthus, Mark van Kleunen, Matthias C. Rillig

## Abstract

- Plants take up nutrients from the soil while investing in absorptive root surface or mycorrhizal partners. Root hairs - a major structure for nutrient uptake and cheap to build - increase the absorptive root surface. As such they are an important component of plant resource economics but largely neglected in root economic concepts so far.
- This is mainly due to data scarcity, which we set out to overcome by measuring root-hair traits on 82 European grassland species in a greenhouse experiment. Using fluorescence and light microscopy, root-hair length and incidence was measured along with mycorrhizal colonization.
- We found a phylogenetically conserved trade-off between plant investment into root hairs and mycorrhiza. A similar trade-off between root-hair incidence and mycorrhiza occurred at the intraspecific level, while patterns were heterogeneous among species. Plant species with high colonization rates showed the highest variability in root-hair incidence.
- We conclude that plants vary along a gradient ranging from investment into root hairs as part of a “do-it-yourself” strategy to collaboration with mycorrhizal fungi while showing intraspecific variation in root-hair incidence. These findings demonstrate that root hairs play a fundamental role in fine-root trait variation and need to be considered when studying belowground plant economic strategies.

## Introduction

The resource economy of plants has been a focal area of studies investigating plant functional traits (Wright *et al*., 2004; Freschet *et al*., 2013a,b; Reich, 2014; Kong *et al*., 2017; Bergmann *et al*., 2020; Weigelt *et al*., 2021, Carmona 2021, Matthus 2025). The general idea is that plants invest carbon in construction and conservation of tissue to ensure the uptake and transport of resources. For aboveground organs - mainly leaves - an economic spectrum of plant strategies has been described and confirmed on a global basis (Wright *et al*., 2004; Reich, 2014; Díaz *et al*., 2016). This spectrum ranges from fast growth and resource acquisition of short-lived organs to slow but steady resource acquisition of organs constructed for longevity. An analogue pattern is found in belowground fine-roots and called the conservation gradient (Bergmann *et al*., 2020; Weigelt *et al*., 2021; Matthus *et al*., 2025).

The concept of a collaboration gradient in root-trait variation that is independent of the conservation gradient and unique to belowground economy has been proposed (Bergmann *et al*., 2020) and found to be a solid pattern across organizational levels and study systems (Matthus *et al*., 2025). This collaboration gradient describes plant strategies in soil exploration ranging gradually from do-it-yourself investment in specific root length (SRL) to outsourcing to mycorrhizal fungal partners with the consequence of larger fine root diameter (AD) and cortex fraction (CF).

Arbuscular mycorrhizal (AM) fungi, which associate with almost 80% of all land plants (Brundrett & Tedersoo, 2018), colonize the root’s cortex and explore the soil with extraradical hyphae (Smith & Read, 2008). AM fungi can take up limiting nutrients like phosphorus and nitrogen, supplying them to the roots in exchange for carbon synthesized by the plant partneŕs aboveground photosynthesis (Bolan *et al*., 1987; Smith & Read, 2008). Besides the exploration of a certain volume of soil, the actual surface and the soil contact of an absorptive plant or fungal structure determines the rate of return on investment of a plant (McCormack & Iversen, 2019). Little is known about traits of fungal extraradical hyphae, but a large body of literature reveals that for the plant itself an effective way to maximize absorptive surface and soil exploration is the production of root hairs (Bhat & Nye, 1973; Gahoonia *et al*., 1997; Bates & Lynch, 2000a,b; Haling *et al*., 2013; Brown *et al*., 2013b). Yet, most likely because of the effort involved in data collection, the coverage of root-hair traits in databases is poor (Iversen *et al*., 2017; Guerrero-Ramirez, 2020; Kattge *et al*., 2020), and their integration into broader plant economics concepts is inconclusive (Zhao *et al*., 2024; Matthus *et al*., 2025).

Root hairs are unicellular epidermal extensions on living fine roots of most land plants (Farquhar, 1996). They enhance nutrient and water uptake of fine roots (Bhat & Nye, 1973; Gilroy & Jones, 2000; Haling *et al*., 2013; Carminati *et al*., 2017; Freschet *et al*., 2021b) contributing to >60% of the plant’s phosphorus-demand (Gahoonia & Nielsen, 1998). Root-hair traits are known to widely vary among species and along environmental gradients of soil fertility (Lambers *et al*., 2008; Holdaway *et al*., 2011), while a large root-hair surface can be realized with long (Yang *et al*., 2015; Haling *et al*., 2016) and/or many (Brown *et al*., 2013a; Marzec *et al*., 2015) root hairs. Furthermore, they have been described to be comparably responsive to soil P availability, in part because of their dynamic growth and life-span (Bates & Lynch, 1996; Zhu *et al*., 2010; Nestler & Wissuwa, 2016). Carbon-cheap, metabolically active (Ma *et al*., 2018) and dynamic in construction compared to fine roots, root hairs might therefore resemble a ‘fast’ economic strategy component within the conservation gradient. So far, no study could provide empirical data to support this link (Matthus *et al*., 2025).

Empirical evidence suggests that the plant species specific beneficial effect of being mycorrhizal is related to root-hair length (Bolan *et al*., 1987; Schweiger *et al*., 1995). Therefore, it has been hypothesized that the carbon investment into root hairs might be an alternative strategy to the mycorrhizal symbiosis on an interspecific (Maherali, 2017) and intraspecific level (Kumar *et al*., 2019) for acquiring soil resources. If this pattern were to be verified for a larger set of species, it would imply that root hairs represent another aspect of a do-it-yourself strategy of plant economics within the collaboration gradient. To date, this hypothesis has not been tested for a larger species set. For four monocotylous families, Betekhtina *et al*. (2023) provide evidence for root hair length to trade-off with AMF colonization levels. Contrary, Guilbeault-Mayers *et al*. (2024) reported a root-hair index (encompassing length and density) to increase with AMF colonization in trees. Parasquive *et al*. (2023) found opposing intraspecific trends of root hair index loadings along the collaboration gradient, depending on tree species. Another study found root hair traits to be independent of the collaboration gradient (Guilbeault-Mayers *et al*., 2024), as has also been proposed by Dallstream & Soper (2024).

It has long been known that non-mycorrhizal plants typically have many and long root hairs while mycorrhizal plants often lack them (Kelley, 1950; Schweiger *et al*., 1995; Jakobsen *et al*., 2005; Brundrett, 2021). Most of these studies only worked with mycorrhizal status as a categorical classification to classify plant functioning. Brundrett & Tedersoo (2018) already noted though, that plant species with many root hairs normally have low mycorrhization and can typically be found in stressful habitats. Still, the current data available originate from different studies conducted under various conditions and measuring different traits and categories, which makes it hard to test for gradual functional trade-offs.

To fill this knowledge gap, we measured root-hair length (HL) and root-hair incidence (HI) as well as mycorrhizal colonization on a large set of grassland species grown under common conditions. We 1) tested for phylogenetic patterns and differences between functional groups and mycorrhizal status. We were interested in whether root hair traits differ between these widely used categories, or whether they gradually change with mycorrhizal colonization. We further aimed to 2) test the hypothesis of an interspecific trade-off between the investment in root hairs and the mycorrhizal partner and to 3) explore the intraspecific variation of root hair traits. Finally, we aimed to 4) integrate interspecific variation of HL and HI into the concept of the root economics space, hypothesizing that root hair investment represents a do-it-yourself strategy additional to the overall increase of SRL.

## Material and Methods

### Species set

The experiment was conducted in the framework of the Biodiversity Exploratories (Fischer *et al*., 2010), a large scale and long term land-use experiment with 150 grassland plots located in three areas in northern, central and southern Germany. From the vegetation records of the Exploratories, we chose a set of 94 grassland species which could be purchased from the commercial seed supplier Rieger-Hofmann GmbH (Blaufelden-Raboldshausen, Germany). This species set encompasses Fabaceae (legumes), non-leguminous dicotyledons (subsequently called forbs) and monocotyledons (subsequently called grasses, but note that Allium schoenoprasum is attached here as a monocotyledonous plant).

### Greenhouse experiment

All data presented here originate from one pot experiment conducted under controlled greenhouse conditions, at the facilities of Freie Universität Berlin, between February and June 2018 (16 h day at 22°C, 8 h night at 15°C). We set up the experiment with 94 initial species, two treatments (with and without mycorrhizal inoculation), and 8 replicates distributed over 4 overlapping time blocks of 6 weeks growing time each. The entire experiment therefore consisted of 4 time blocks x 94 species x 2 treatments x 2 replicates = 1504 experimental units. Whenever a replicate did not survive, we tried to substitute it in the next time block. Nevertheless, some species did not reach a replication of 8 per treatment and some did not germinate at all. In the final analysis, we only included species with a minimum of 3 successful replicates per treatment leading to a total of 1151 experimental units of 82 species from 20 families.

Prior to the first time block, all seeds were surface sterilized in paper tea bags for 3 min in 7% bleach followed by washing in de-ionized (DI) water until the smell of bleach was gone. The seeds were dried at 20°C and stored until sowing for subsequent time blocks. Seeds germinated in plastic boxes filled with 1:1 steamed sand and vermiculite (1-3 mm, ISOLA Vermiculite GmbH; Sprockhövel, Germany). Based on germination times recorded in pre-experiments, we sowed the seeds to assure that all seedlings were in the cotyledon stage or had their first leaves developed at time of transplanting. Seedlings were transplanted into plastic cones (410 ml 0.41 L; Stueve & Sons; USA) filled with the same substrate as for germination.

The mycorrhizal treatment was realized as follows: after filling the cone to c. ¾ we added a 30 ml horizon of a 1:1 mixture of steamed sand and mycorrhizal inoculum in vermiculite (INOQ Agri, Inoq GmbH, Schnega, Germany). According to the supplier, the inoculum contains 145 spores/ml of *Rhizophagus irregularis* propagated on vermiculite (1-2 mm) under non-sterile greenhouse conditions. *Rhizophagus irregularis* is a generalist AM fungus associating with almost all mycorrhizal plants (van der Heijden *et al*., 2015). To account for other soil microbes that might be present in AM inoculum produced under non-sterile conditions, we prepared a microbial wash from the *Rhizophagus* inoculum (20 µm mesh, soil:water-ratio: 1:2) for the non-mycorrhizal treatment. We carefully adjusted the amount of inoculum as well as the amount of DI water used to prepare the microbial wash to make sure that each pot received the approximate same number of microbial units irrespective of the treatment. To control for nutrients and physical structure of the AM inoculum, we autoclaved the solid inoculum used to prepare the microbial wash and added a 1:1 mixture with steamed sand as a horizon to the non-mycorrhizal treatment. For both treatments the added horizons were covered with another layer of ∼30 ml substrate to avoid cross contamination. During transplanting, seedlings of the non-mycorrhizal treatment received 30 ml of the microbial wash, while seedlings of the mycorrhizal treatment received 30 ml of DI water. We replaced seedlings that died shortly after transplanting during the first week.

Within each time block, plants grew for 6 weeks in the cones before harvest. All cones were fully randomized at time of transplanting and were rearranged every two weeks. Plants received 25 ml of DI water 3 times a week; two weeks and four weeks after transplanting, they received 25 ml of a ¼ strength Hoagland solution (recipe available in Lachaise *et al*., 2021) instead.

At time of harvest, aboveground and belowground biomass of the plants were separated. Roots were first rinsed with water. Three first order roots per plant were carefully cut, transferred to 10% formalin at pH 7 (ROTI Histofix, Carl Roth, Karlsruhe, Germany) in Phosphate Buffered Saline (PBS) buffer and kept at 4 °C for overnight fixation. The next day, the formalin solution was first replaced by PBS buffer twice for approx. 2 hours each and finally by a solution of 70% ethanol, 5% glycerin and 25% DI water for long term preservation. The remaining roots were carefully washed by hand during harvest, transferred to cold DI water and kept at 4°C. Within a week they were scanned in water-filled plastic trays using an Epson perfection 800 Photo scanner at a resolution of 800 dpi. As root systems were small and young, we decided to measure traits on the entire root system. This included mainly first to third order roots and a small fraction of higher order roots. 99.98 % of root length within the entire experiment belonged to roots with a diameter < 2 mm.

### Trait measurements

Total root length and volume as well as the average root diameter (D [mm]) were measured using WinRhizo 2017 software (Regent Instruments Inc., Québec, Canada). Aboveground and belowground dry biomass was determined after drying at 60°C for at least three days. The mycorrhizal growth response (MGR) of each plant species was calculated as MGR= ln[total dry biomass inoculated / total dry biomass non-inoculated] (Hoeksema *et al*., 2010; Maherali, 2014). All other root traits were measured within the mycorrhizal treatment assuming that this resembles the natural soil biotic condition. Root dry biomass was used to calculate the specific root length (SRL – root length/dry biomass [m/g]) and root tissue density (RTD – dry biomass/root volume [g/cm³]) by calculating the volume as the sum of 0.2 mm diameter size classes according to Rose (2017).

Root-hair length (HL [µm]), cortex fraction (CF [%]) and first order root diameter (D_first_ [mm]) were measured on the preserved first order root tips of three randomly chosen replicates per species from the mycorrhizal treatment using a fluorescence microscope (Zeiss Axio Imager 2, Carl Zeiss AG, Oberkochen, Germany). One root per replicate was randomly picked and dyed in 0.01% Calcofluor-white (Thermo Fisher Scientific, Waltham, USA) for 5-10 seconds. Subsequently, it was rinsed in DI water for a few seconds, mounted on a slide and carefully covered with a cover slip without applying pressure. As Calcofluor-white binds to cellulose, it helps distinguish plant cell walls including those of fine root hairs. As all roots were small and translucent there was no need for cross sectioning to measure stele and cortex diameter (Fig. S1). Microscopic images were taken with a Zeiss AxioCam at a magnification of x50 using a 430 nm fluorescence filter. For each replicate, several images were taken using the functions “Z-Stacks” and “Tiles” to display a continuous segment of 5 mm within the mature root hair zone. The “Tiles” function merges several images along the root while the “Z-Stacks” function combines images vertically, thereby producing an in-focus image throughout the entire range of depths. We defined the Z-Stacks to range from the middle of the stele to the upper epidermal layer of the root just beneath the cover slip. The first order root diameter (D_first_) as well as the stele diameter were measured at three positions along the image. We calculated the cortex fraction (CF) as the percent area of a first order root cross section that is occupied by tissue outside the stele (Freschet *et al*., 2021a). Mean values of D_first_ and CF were first calculated at replicate level and subsequently at the species level. HL was measured according to Delhaize *et al*. (2012). In brief, we divided the 5 mm root segments into 5 sub-segments of 1 mm each and measured the length of the longest root hair on each side of the root in each sub-segment (i.e. 2 root hairs per sub-segment). All 10 measurements per 5 mm root segment were averaged to calculate the mean HL per replicate. Mycorrhizal status (obligate mycorrhizal, facultative mycorrhizal, non-mycorrhizal) was assigned on species level according to the FungalRoot database (Soudzilovskaia *et al*., 2020). In case of conflicting status reports within the database, we followed the provided expert recommendations.

For the determination of the percentage of mycorrhizal colonization (%M) and the root-hair incidence (HI [%]), we used representative subsamples of the dried root systems of the three replicates from the mycorrhizal treatment of each species (Freschet *et al*., 2021a). Furthermore, one replicate per species in the non-mycorrhizal treatment was randomly chosen and checked for AM colonization. Roots were first cleared in 10% (w/v) KOH for 15 min at 80°C and then stained in 0.05% (w/v) Trypan Blue in lactoglycerol for another 15 min at 80°C. Mycorrhizal colonization was determined with the magnified intersection method (McGonigle *et al*., 1990) at a magnification of x200, using a minimum of 30 root pieces on a slide to count presence or absence of mycorrhizal structures in general (colonization rate) and of arbuscules in specific (rate of arbuscular colonization) in 50-100 intersects. Mycorrhizal hyphae were identified based on their staining, their growing habit (intraradical between cortical cells) and their missing of irregular septation. Due to the commercial inoculum, there was very little contamination with other fungi. For the non-mycorrhizal treatment, mycorrhizal colonization rates between 1% and 6% were detected for 6 out of 82 replicates suggesting very limited contamination. Within the mycorrhizal treatment colonization rates of up to 86% and rates of arbuscular colonization up to 72% clearly confirmed a successful inoculation. HI was determined simultaneously and analogously to %M, recorded as presence or absence at each intersect (Siqueira & Saggin-Júnior, 2001), giving a proxy of how much percent of the root length was covered by root hairs.

The coefficients of variation of HI and HL (cvHI, cvHL) were calculated by using the general R function cv(x) that computes the sample coefficient of variation as (SD/mean)*100. To display the within-species correlation between %M and HI of all species while accounting for overall between-species differences in both traits, we normalized the data by coding the intraspecific median of the three trait records per species as 0, the lower value as = lower value – median value and the higher value as = higher value – median value.

Root nitrogen concentration (N [%]) was measured on the three replicates, after drying and milling the roots, using an Elemental Analyzer (Euro EA, HEKAtech, Wegberg, Germany).

### Analysis

All analyses were carried out in R version 3.6.3 (R Core Team, 2020). To explore phylogenetic patterns in the root hair data we used the function drop.tip() from the package ape (Paradis *et al*., 2004) to prune the DaPhnE phylogeny (Durka & Michalski, 2012) for our species set and the function phylosig() as well as phylo.heatmap() from the package phytools (Revell, 2012) to calculate the phylogenetic signal of all traits and to display trait variation along the tree by using color palettes from the package viridis (Garnier, 2018). The package ggplot2 (Wickham, 2010) was used to display violin plots, the pairwise correlation heatmap and the correlation between %M and HI at intraspecific level and the package cowplot (Wilke, 2024) was used for multipanel figures.

Prior to the calculation of pairwise correlations and principal component analyses (PCA), we improved data distribution by applying log transformation for all traits except CF, HI and %M, which we transformed using the function logit() from the gtools (Warnes *et al*., 2020) package, since these traits varied between 0 and 1. The function rcorr() from the package Hmisc (Harrell, 2020) was used to calculate Pearson’s correlations of all trait pairs, and the functions comparative.data() and pgls() from the package caper (Orme *et al*., 2018) were used to calculate phylogenetically corrected pairwise correlations. We determined the phylogenetically corrected correlation coefficient by taking the square root from the adjusted *r*² of the model multiplied by -1 in case of a negative regression coefficient while assigning *r*=0 in case of negative adjusted *r*² values. The phylogenetically informed PCAs were calculated using phyl.pca() from the package phytools (Revell, 2012) and displayed using functions from the package shape (Soetaert, 2014). Ellipsoids were plotted using the package ellipse (Murdoch & Chow, 2024). Permanova based on euclidean pairwise distances in PCA space among groups were performed using the pairwise.adonis() function from the package pairwiseAdonis (Martinez Arbizu, 2019).

We further used the packages dplyr (Wickham *et al*., 2020a), Rmisc (Hope, 2013), raster (Hijmans, 2020), data.table (Dowle & Srinivasan, 2020) and devtools (Wickham *et al*., 2020b) for general data handling and exploration.

## Results

### Root hair traits show a phylogenetically conserved pattern

Root-hair length and incidence showed strong phylogenetic signals (Fig. 1, Table S1) and were highly positively correlated (Fig. S2). Monocotyledons showed many long root hairs, with the hairless *Allium schoenoprasum* being the only exception, while legumes (Fabaceae) had few and short root hairs (Fig. 1). Within the other dicotyledonous families, Asteraceae showed low values while Polygonaceae, Caryophyllaceae and Brassicaceae showed high values for both root-hair length and incidence. Throughout the entire set of species, mycorrhizal colonization showed a completely inverted pattern which was strongly phylogenetically conserved as well (Fig. 1, Table S1). Clades with long root hairs and high root-hair incidence were poorly colonized by mycorrhizal fungi, while clades with short and few root hairs showed high colonization rates. Root-hair length and incidence were both negatively correlated with mycorrhizal colonization. This pattern disappeared for root-hair length after phylogenetic correction (Fig. S2).

**Fig. 1:**
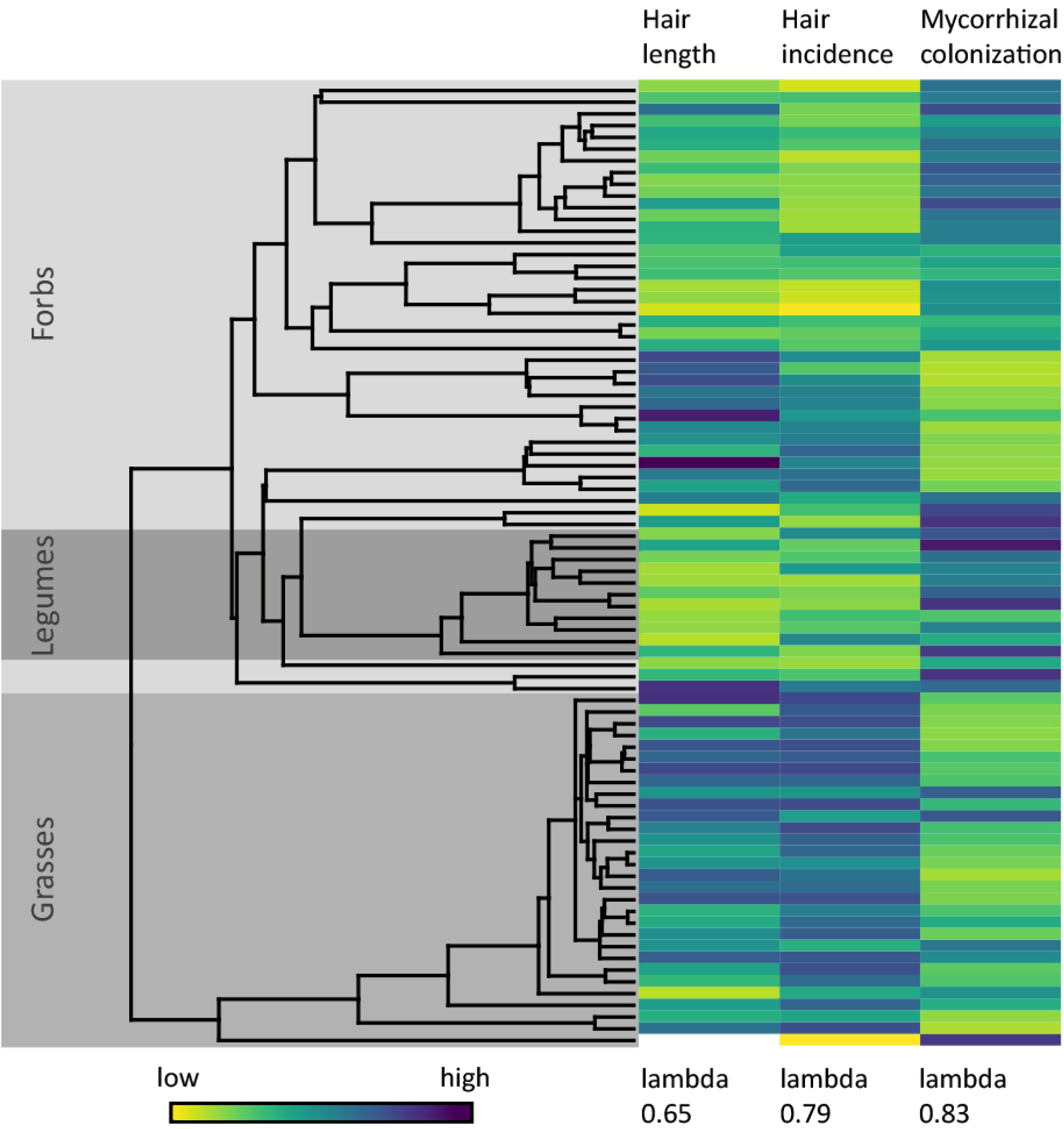
The phylogenetically conserved trade-off between the investment in root hairs and mycorrhization. Colour-coded are the species mean values of root-hair length and incidence as well as percent mycorrhizal colonization to the right and the corresponding phylogenetic tree with broader taxonomic groups to the left. Trait values are standardized to the same range, colour-coded from yellow (low) via green (medium) to blue (high). Phylogenetic signal of each trait is displayed as Pageĺs lambda. Grasses are shaded in medium grey, non-leguminous forbs in light grey and leguminous forbs in dark grey. *Allium schoenoprasum* L. with the lowest hair incidence had no root hairs in the samples for determination of hair length leading to a missing value.

Examining plant functional types, root-hair length was lower in legumes than in grasses and forbs, with grasses having the longest root hairs overall (Fig. 2a). The coefficient of variation in root-hair length did not differ significantly among these plant functional groups, even though grasses tended to have the least variation (Fig. 2e). Root-hair incidence was higher and its coefficient of variation was lower in grasses than in both legumes and forbs (Fig. 2b,f).

**Fig. 2:**
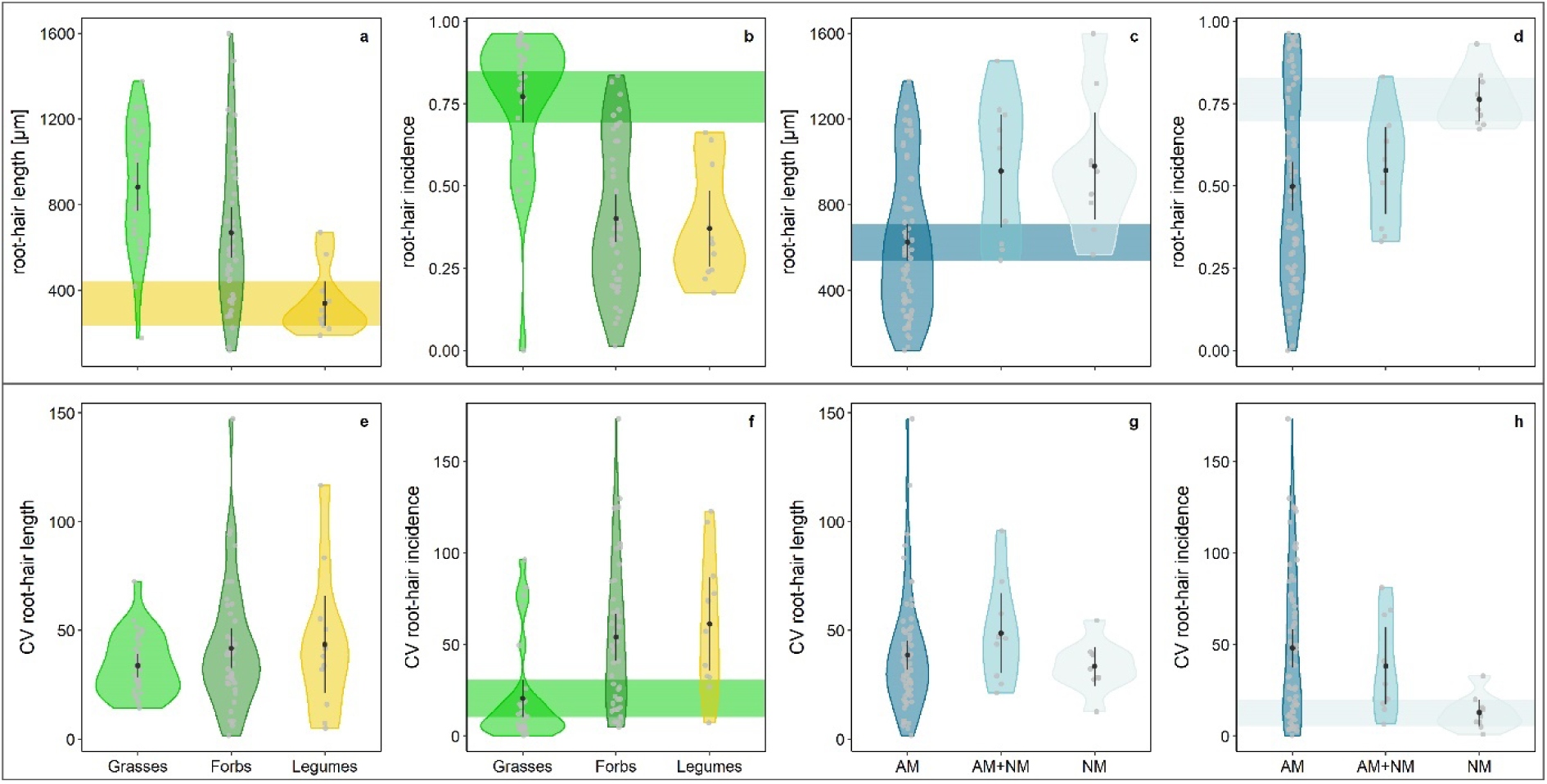
Variation in root hair traits according to plant functional type and mycorrhizal status. Displayed are raw data (upper panels) of species mean root-hair length (HL, panel a, c) and incidence (HI, panel b, d) as well as the coefficient of variation (lower panels) in root-hair length (cvHL, panel e, g) and incidence (cvHI, panel f, h). Displayed are kernel density distributions and group means (black dots) with 95% confidence intervals. Non-overlapping confidence intervals are highlighted by a coloured ribbon to visualize group differences. Plant functional types: grasses, forbs, legumes; mycorrhizal status: obligate mycorrhizal (AM), facultative mycorrhizal (AM-NM), non-mycorrhizal (NM).

### Mycorrhizal status is not a strong predictor of root-hair traits

Of the 82 grassland species tested in the experiment, 64 were obligate mycorrhizal, 9 facultative mycorrhizal and 9 non-mycorrhizal, as classified according to the FungalRoot database. Mycorrhizal status did not predict species root-hair length or its coefficient of variation well, even though obligate mycorrhizal species tended to have shorter root hairs (Fig. 2c). Root-hair incidence instead was lower in obligate mycorrhizal species than in non-mycorrhizal species, while facultative mycorrhizal species had intermediate values (Fig. 2d). The coefficient of variation of root-hair incidence showed the opposite pattern with obligate mycorrhizal species being more variable than non-mycorrhizal species, and facultative mycorrhizal species showing intermediate values again (Fig. 2h).

### An ecological trade-off between root hairs and mycorrhization

The phylogenetically informed PCA revealed a strong trade-off (PC 1 = 45%) between high root-hair incidence and length on one end, and high mycorrhizal colonization rates on the other, accompanied by an increase in variation in root-hair incidence (Fig. 3, Table S2). PC 2 explained 23% of variation, with the coefficient of variation of root-hair length influencing this axis most strongly.

**Fig. 3:**
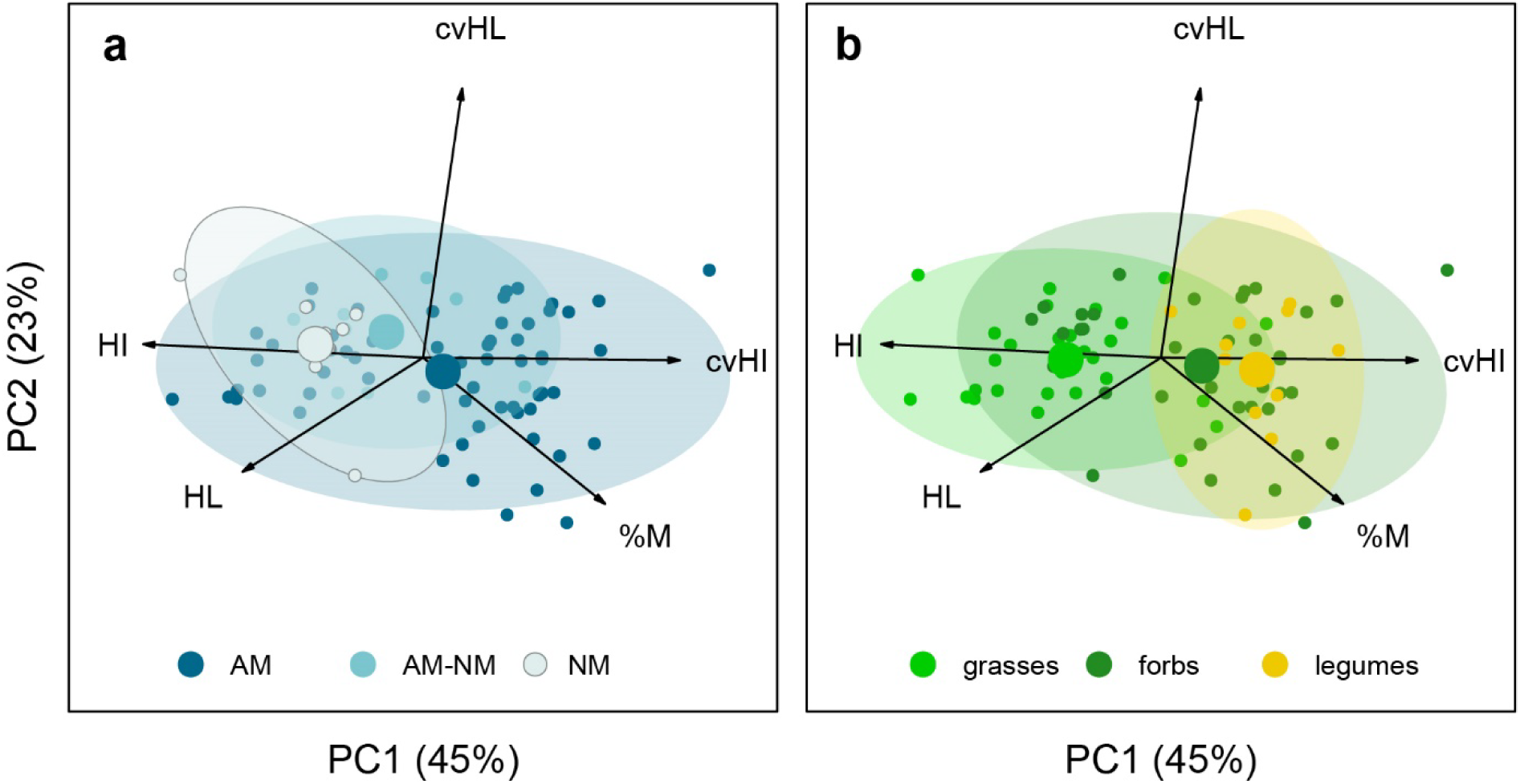
Phylogenetically informed principal component analysis of root hair traits and mycorrhizal colonization rate. Panel a displays species based on their mycorrhizal status (obligate mycorrhizal - AM, facultative mycorrhizal - AM-NM, non-mycorrhizal - NM) while panel b displays species based on their plant functional group (grasses, forbs, legumes). Ellipsoids and large dots display 95% confidence intervals and centroids. PCA results can be found in table S2. HL – hair length, HI – hair incidence, cvHL – coefficient of variation in hair length, cvHI – coefficient of variation in hair incidence, %M - percent mycorrhizal colonization.

Mycorrhizal status as well as plant functional types affected species locations within the PCA (Table S3). Non-mycorrhizal species differed from obligate mycorrhizal species by being closely aggregated at high values of root-hair incidence on PC 1. Grasses differed from both forbs and legumes by showing high root-hair incidence as well, even though considerable variation occurred within each of the functional types. Legumes were located at high values of mycorrhizal colonization and variation in root-hair incidence on PC 1 while spanning the entire range of PC 2.

### At the intraspecific level, root-hair incidence correlates with mycorrhizal colonization rate but species show strong heterogeneity

Despite low within-species replication (n = 3 individual plants per species scored for root hair traits), we could detect an overall association between root-hair incidence and mycorrhizal colonization rate (Fig. 4). Individual plants with higher colonization rates had lower root-hair incidence than less-colonized individuals within the same species. Overall, we found a slight negative correlation (slope = -0.29, p<0.001) with small confidence intervals. However, there was considerable variation among species. No intraspecific correlation was found between the mycorrhizal colonization rate and root-hair length.

**Fig. 4:**
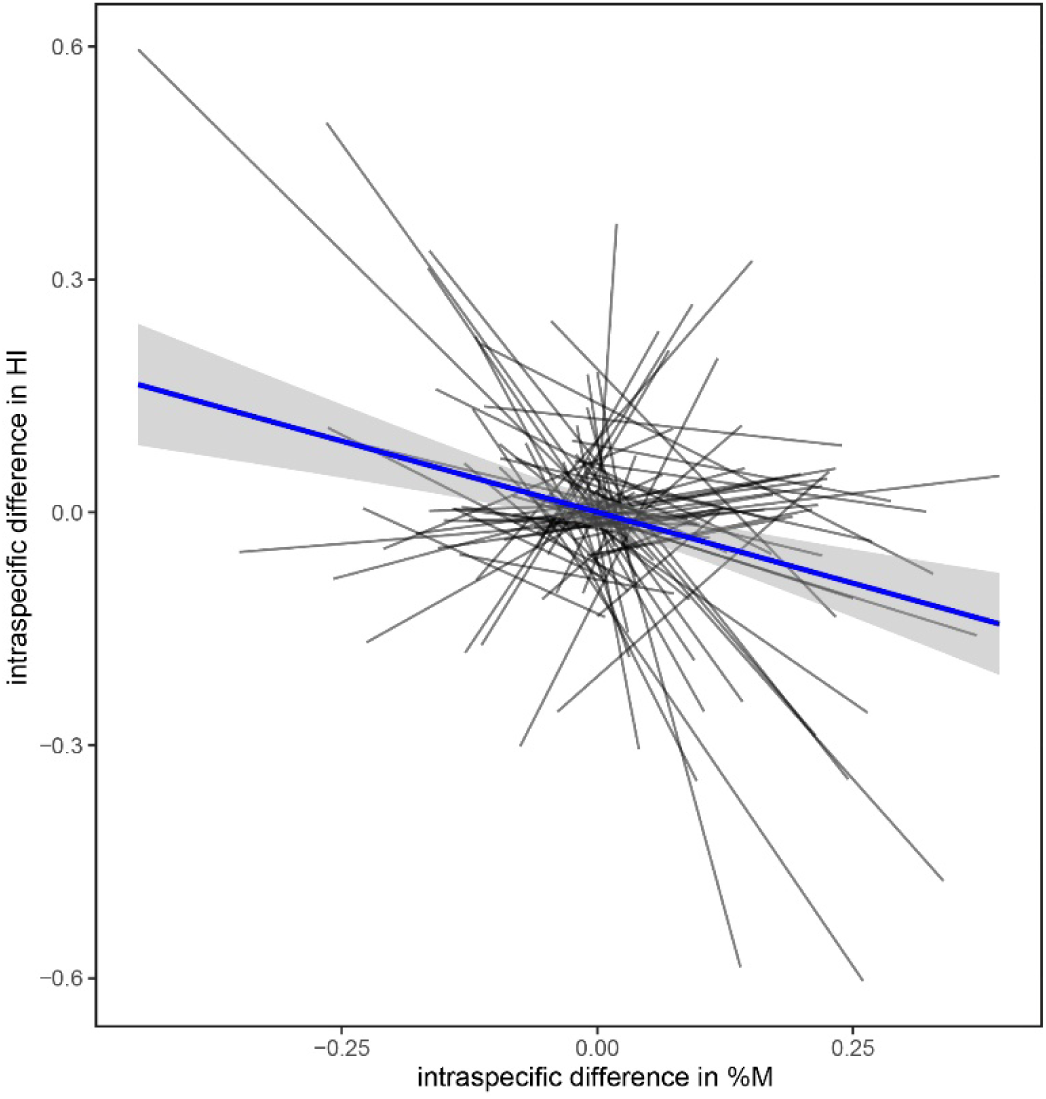
Intraspecific correlation of root-hair incidence and mycorrhizal colonization. Displayed is the relative difference in mycorrhizal colonization (%M) and root-hair incidence (HI) within each species as well as the overall correlation with 95% confidence interval.

### Root-hair traits add to the root economics space

The inclusion of root-hair traits introduced a new dimension to the root economics space. The first axis (22%) of the extended PCA (Fig. 5, Fig. S3, Table S4) was dominated by specific root length on one end and root average diameter and diameter of first order roots on the other, accompanied by variation in cortex fraction but also root-tissue density. The trade-off between the investment in root-hair length and incidence on one end and the coefficient of variation of root-hair incidence on the other dominated the second axis (20%). Mycorrhizal colonization intensity and mycorrhizal growth response were associated with variation in root-hair incidence. Root-tissue density loaded strongest on the third axis (14%) together with the coefficient of variation of root-hair length and antagonistically to root-nitrogen concentration and cortex fraction.

**Fig. 5:**
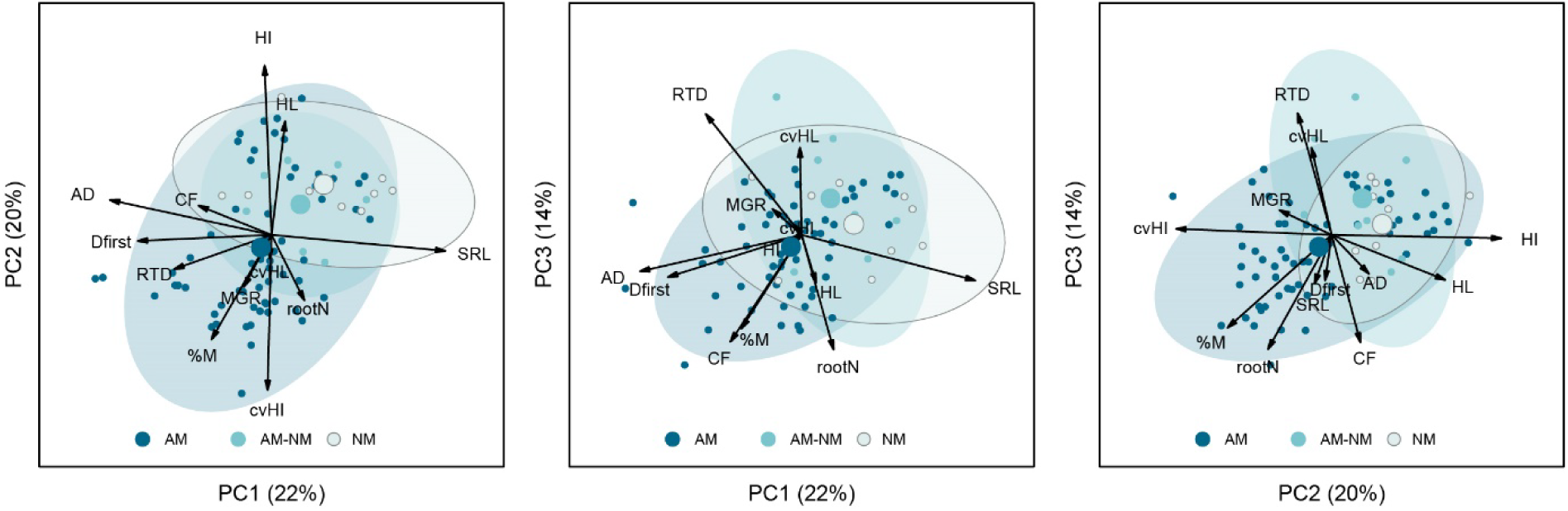
Extended phylogenetically informed principal component analysis. Displayed are species based on their mycorrhizal type (obligate mycorrhizal - AM, facultative mycorrhizal – AM-NM, non-mycorrhizal - NM). Ellipsoids and large dots display 95% confidence intervals and centroids. PCA results can be found in Table S4. HL – hair length, HI – hair incidence, cvHL – coefficient of variation in hair length, cvHI – coefficient of variation in hair incidence, %M - % mycorrhizal colonization, SRL – specific root length, AD – average diameter, D_first_ – diameter of first order roots, CF – cortex fraction, RTD – root tissue density, N – root nitrogen concentration, MGR – mycorrhizal growth response.

Plants of different mycorrhizal status as well as different plant functional types differed in their root economic strategies within the space (Table S5). Non-mycorrhizal and facultative mycorrhizal plants differed from obligate mycorrhizal plants, while non-mycorrhizal plants showed the highest specific root length on PC1 and highest root-hair length and incidence on PC2, and there was no general pattern along PC3. Obligate mycorrhizal plants spanned the entire space but clearly showed the highest values for root diameter on PC1, lowest root-hair incidence and highest colonization rate on PC2 and highest root-nitrogen concentration on PC3. Legumes were located at high root diameter, cortex fraction and colonization rate as well as high root-nitrogen concentration. Grasses showed a clear trend towards high root-hair length and incidence as well as specific root length. As such, grasses and legumes formed distinct groups almost without overlap, while forbs spanned the entire root economics space.

## Discussion

### Plants can either invest in root hairs or rely on mycorrhizal partners while maintaining variation in root-hair incidence

We found a striking pattern of an evolutionarily conserved trade-off between plant investment in root hairs – specifically their incidence – and mycorrhizal symbiosis. The phylogenetic conservation occurred at high taxonomic levels; grasses showing high root-hair incidence and length, paired with low mycorrhizal colonization rates, while legumes exhibited the opposite pattern (Fig. 1 and 2). This supports existing knowledge on grasses and legumes (Hill *et al*., 2006). Non-leguminous forbs exhibited a range of strategies along the entire gradient of variation. This is an expectable result, given that forbs are not monophyletic and comprise all forms of mycorrhizal status. An evolutionarily deep-rooted phylogenetic signal might be the reason why many correlations between raw trait data disappeared after phylogenetic correction.

In contrast to the level of mycorrhizal colonization, we found mycorrhizal status to be a weak predictor of root-hair traits. The traditional mycorrhizal status classification of species as being obligate, facultative or non-mycorrhizal and the respective definitions have been discussed lately (Cosme *et al*., 2018; Brundrett & Tedersoo, 2019). Cosme *et al*. (2018) argue that species classified as being non-mycorrhizal can have low levels of colonization and even a few arbuscules. We found the same pattern in our species classified as non-mycorrhizal with low colonization rates and no or very few arbuscules. Moreover, we found that these species had high root-hair length and incidence, hence resembling an extreme do-it-yourself trait syndrome.

Non-mycorrhizal plants can be subdivided based on their phosphorus (P) acquisition strategy as P-scavengers, which rely on dissolved P, and P-miners, which exude organic compounds that release fixed P (Lambers *et al*., 2008; Lambers & Teste, 2013; Yu *et al*., 2020). *Carex vulpina*, as the only non-mycorrhizal species with a P-mining strategy in our dataset showed the highest HI but only an average HL within the non-mycorrhizal status. A larger number of Proteaceae type species (P-miners) would be needed to draw general conclusions about HL and HI patterns of the different non-mycorrhizal P acquisitions strategies. Hence, our results can be considered representative only for P scavenging species.

The classification of facultative mycorrhizal species includes both species that are mycorrhizal only under specific circumstances (like nutrient deficiencies) and species that always have low colonization rates; hence plants with different ecological strategies (Brundrett & Tedersoo, 2019; Soudzilovskaia *et al*., 2020). Furthermore, the status ‘facultative’ can be misleading in case of conflicting observations for species with an overall low data record. Accordingly, we found strong overlap in root-hair traits between obligate and facultative mycorrhizal species, even though the latter tended to have more and longer root hairs as we would have expected given the fact that they have lower colonization rates (Fig. S4). The categories of facultative and obligate mycorrhizal species seem to not be informative for root hair patterns. However, non-mycorrhizal plants differed strongly from obligate mycorrhizal plants by having higher root-hair incidence and lower variation therein. This pattern also dominated the overall gradual trade-off between the investment in root hairs and mycorrhiza: a strong investment in root-hair incidence was accompanied by low variation of the same trait. Species with high mycorrhizal colonization rates produce fewer root hairs but encompass more intraspecific variation. The coefficient of variation provided us with a scale-independent measure of variation. It should be noted though that at a given standard deviation, it is inversely related to the mean value, hence mathematically favoring high mean trait values to coincide with low variation of the same trait. Given the fact that being mycorrhizal can be a competitive advantage in many though not all terrestrial habitats (Brundrett & Tedersoo, 2018), it remains to be studied how root hair traits add to the filtering of environmental variation for species occurrence (Laughlin *et al*., 2021).

### Intraspecific variation in root-hair incidence and mycorrhizal colonization mirrors the interspecific pattern

Although this experiment was not designed to test for intraspecific variation, we could show that overall, the within-species root-hair incidence was higher at lower colonization rates. We cannot determine if this variation originates from a plastic response of the plant to different colonization levels of the AM fungus or from genetic variation between plant individuals. Further research is needed to evaluate this question and to determine cause and effect. Plasticity in both root-hair length and incidence has been reported in response to soil P (Bates & Lynch, 1996, 2000b; Zhu *et al*., 2010) as well as mycorrhizal inoculation (Price *et al*., 1989; Sun & Tang, 2013; Wu *et al*., 2016). Suggestions about the resource costs of root hairs being higher (Price *et al*., 1989) or lower (Brown *et al*., 2013a) than those of the mycorrhizal symbiosis differ widely, while soil moisture and P availability (Brown *et al*., 2013a; Fort *et al*., 2015; Ma *et al*., 2021) further mediate the effects. Given the design of our study with homogeneous soil fertilization, we cannot test the effect of soil P on root hair traits and their variability. Specifically, it remains to be studied if facultative mycorrhizal species show higher coefficients of variation in root-hair incidence and length under different levels of soil P. Furthermore, patterns might change over time, given the fact that we analyzed young plants. Nevertheless, our results suggest an overall intraspecific trade-off between root-hair incidence and mycorrhizal colonization mirroring the interspecific pattern and leading to stronger variation in root-hair incidence in obligate mycorrhizal species with high colonization rates.

### Root hairs add to the do-it-yourself strategy of plants

The trade-off between root hairs and mycorrhizal colonization rate defined the second axis of the principal component analysis on all traits, with the root traits of the collaboration gradient dominating the first and those of the conservation gradient the third axis. The first axis resembled the collaboration gradient with a trade-off between ‘do-it-yourself’ with high SRL and ‘outsourcing’ with high root diameter and cortex fraction as expected within the framework of the root economics space (Bergmann *et al*., 2020; Ding *et al*., 2020; Wen *et al*., 2022). RTD also loaded on axis 1 – though less than on axis 3 - with a considerable amount of variation, being negatively correlated to SRL. This correlation has been reported before (Eissenstat, 1992; Reich, 2014) and might originate from the fact that, for a given diameter SRL, has to increase with decreasing RTD (Ostonen *et al*., 2007). This might be the most important driver behind former detection of a one dimensional root economics spectrum that parallels leaf economics (Freschet *et al*., 2010; Reich, 2014).

On axis 2, root-hair length and incidence behaved antagonistically to the degree of variation in root-hair incidence accompanied by mycorrhizal colonization rate and growth response. With 20% variance, the trade-off explained a considerable amount of variation within the entire trait space. Mycorrhizal colonization was less strongly associated with the first axis than with the root-hair dominated second axis, though the bivariate correlation with cortex fraction was strong as expected in the concept of the collaboration gradient. Since mycorrhizal colonization was measured on the same microscopy slides as root-hair incidence, while root-hair length and cortex fraction were measured separately, we do not expect a methodological bias here. As for the correlation with %M, both traits of the first and the second axis link to the functional concept of collaboration. This is in line with the scheme proposed by Wen *et al*. (2019) who found species to either rely on a root morphology of high absorptive surface, which can be achieved in different ways (in their case by high SRL or branching) or on mycorrhizal symbiosis and a high root diameter for P-scavenging. Root hairs, as another absorptive structure added to the concept, are also involved in P-mining by exudation (Wen *et al*., 2022) but the respective impact on their association with the collaboration gradient is yet to be explored as P-mobilizing exudates were not measured in the current study. Taken together, it remains to be tested on a larger set of species, if the inclusion of root hair traits widens the collaboration gradient to a plane encompassing different strategies for do-it-yourself resource acquisition.

The third axis resembled the conservation gradient proposed as the belowground analogue of the fast-slow economic spectrum in leaves (Weigelt *et al*., 2021), with RTD representing the ‘slow’ and root-N concentration representing the ‘fast’ strategy. The degree of variation in root-hair length was also associated with the ‘slow’ strategy on axis three. We can only speculate that this might be related to the fact that slow growing species invest in fine roots with a longer lifespan. Hence those species might keep the ability to alter the length of the comparably more short-lived root hairs given the fact that their surface provides the main absorptive structure for those species (Fort *et al*., 2015). Cortex fraction loaded on the ‘fast’ side of axis three. We hypothesize that this unexpected link might occur because AMF also enhance species N uptake under limiting conditions (Govindarajulu *et al*., 2005; Hodge & Fitter, 2010). As root-N concentration, cortex fraction and colonization rate were measured on the same replicates, the effect of mycorrhizal colonization rate on root-N concentration and a resulting positive correlation (r=0.3) might be overestimated by our data, which were measured on plants growing under relatively low nutrient conditions. Furthermore, our experiment was restricted to a single AMF species and excluded plant mycorrhizal types other than arbuscular mycorrhiza. Ectomycorrhizal species tend to occupy areas of the ‘slow’ strategy in the root economics space (Bergmann *et al*., 2020), hence adding variation to the conservation gradient that is not covered in the present experiment. It is also important to notice that, due to the nature of the ectomycorrhizal symbiosis with fungal hyphae covering entire fine roots and leading to fast degradation of root hairs (Farquhar, 1996) the importance of root-hair traits might change in a global dataset.

## Conclusions

We can support the hypothesis that investment into root hairs and mycorrhizal partnerships are alternative ecological strategies for soil exploration and resource uptake with a strong evolutionary history. This interspecific ecological trade-off is mirrored at the intraspecific level with plants showing more root hairs at lower mycorrhizal colonization rates. Strong heterogeneity between species calls for further investigations of intraspecific patterns. A high degree of variation in root-hair incidence is associated with high mycorrhizal colonization rates and growth response at the species level. The ecological trade-off between the investment in root hairs and the degree of variation in incidence being intraspecifically correlated with mycorrhizal colonization rates dominates the second axis of the root economics space. We conclude that variation in root-hair patterns is neither fully aligned with the conservation gradient nor the existing concept of the collaboration gradient but rather introduces a new dimension of variation into the picture. Still, regarding the strong trade-off with mycorrhizal colonization, we consider root hairs, and specifically their incidence, to add to the ecological strategy of ‘do-it-yourself’. Hence, we find the concept of collaboration to span the first and second and the conservation gradient to represent the third axis of variation in the root economics space. These results present strong evidence that root hairs are a considerable source of variation in fine root morphology that should be considered when studying belowground plant functioning.

## Acknowledgements

We thank Erik Faltin, Cathrin Schierenbeck, Anja Wulf, Max Fussan, Maxi Bergmann and many others for help with root washing and scanning. We further thank Julien Bachelier for use of the fluorescence microscope and in particular Maria Schauer for sharing her knowledge about fixation and microscopy of plant material. We also thank the managers of the three Biodiversity Exploratories, Konstanz Wells, Swen Renner, Kirsten Reichel-Jung, Sonja Gockel, Kerstin Wiesner, Katrin Lorenzen, Andreas Hemp, Martin Gorke and Miriam Teuscher, and all former managers for their work in maintaining the plot and project infrastructure; Christiane Fischer for giving support through the central office, Andreas Ostrowski for managing the central data base, and Markus Fischer, Eduard Linsenmair, Dominik Hessenmöller, Daniel Prati, Ingo Schöning, François Buscot, Ernst-Detlef Schulze, Wolfgang W. Weisser and the late Elisabeth Kalko for their role in setting up the Biodiversity Exploratories project. The work has been funded by the DFG Priority Program 1374 “Infrastructure-Biodiversity-Exploratories”. Field work permits were issued by the responsible state environmental offices of Baden-Württemberg, Thüringen, and Brandenburg. We acknowledge funding from the German Research Foundation (DFG, grants 432975993 to JB, KL 1866/12-1 to MvK and 323522591 to MR).

## Author contribution

JB designed and performed the experiment, ran the analyses and wrote the paper. TL contributed to the analysis and the conceptual development of the study and revised the paper. KB and EB participated in the experiment and data exploration. EM revised the paper. MvK and MR contributed to the study design and revised the paper.

## Data availability

This work is based on data elaborated by the RootFun project (323522591) and further analyzed within the HAIRphae project (432975993) of the Biodiversity Exploratories program (DFG Priority Program 1374). All data used is publicly available in the Biodiversity Exploratories Information System (http://doi.org/10.17616/R32P9Q) under dataset ID ##### (will be linked upon acceptance) or in databases referenced. The authors explicitly encourage appropriate usage and database implementation of the data.

## Supporting information

**Fig. S1:** Microscopic image of first order root of *Helictrotrichon pubescens* (HUDS.) DOMORT., Poaceae, obligate mycorrhizal.

**Fig. S2:** Pairwise correlations of all measured traits.

**Fig. S3:** Extended phylogenetically informed principal component analysis.

**Fig. S4:** Variation in mycorrhizal colonization (%M) between plants of different mycorrhizal status.

**Table S1:** Phylogenetic signal of all measured traits.

**Table S2:** Phylogenetically informed principal component analysis of root hair traits and %M.

**Table S3:** Permanova based on pairwise dissimilarities of plant functional types and mycorrhizal types within the principal component analysis displayed in Table S2.

**Table S4:** Extended phylogenetically informed principal component analysis.

**Table S5:** Permanova based on pairwise dissimilarities of plant functional types and mycorrhizal types within the extended principal component analysis displayed in Table S4.

## References

Bates TR, Lynch JP. 1996. Stimulation of root hair elongation in Arabidopsis thaliana by low phosphorus availability. Plant, Cell and Environment 19: 529–538.

Bates TR, Lynch JP. 2000a. The efficiency of Arabidopsis thaliana (Brassicaceae) root hairs in phosphorus acquisition. American Journal of Botany 87: 964–970.

Bates TR, Lynch JP. 2000b. Plant growth and phosphorus accumulation of wild type and two root hair mutants of Arabidopsis thaliana (Brassicaceae). American Journal of Botany 87: 958–963.

Bergmann J, Weigelt A, Van der Plas F, Laughlin DC, Kuyper TW, Guerrero-Ramirez N, Valverde-Barrantes OJ, Bruelheide H, Freschet GT, Iversen CM, et al. 2020. The fungal collaboration gradient dominates the root economics space in plants. Science Advances 6: 1–9.

Betekhtina AA, Tukova DE, Veselkin D V. 2023. Root structure syndromes of four families of monocots in the Middle Urals. Plant Diversity 45: 722–731.

Bhat KKS, Nye PH. 1973. Diffusion of phosphate to plant roots in soil - I. Quantitative autoradiography of the depletion zone. Plant and Soil 38: 161–175.

Bolan NS, Robson AD, Barrow NJ. 1987. Effects of vesicular-arbuscular mycorrhiza on the availability of iron phosphates to plants. Plant and Soil 99: 401–410.

Brown LK, George TS, Barrett GE, Hubbard SF, White PJ. 2013a. Interactions between root hair length and arbuscular mycorrhizal colonisation in phosphorus deficient barley (Hordeum vulgare). Plant and Soil 372: 195–205.

Brown LK, George TS, Dupuy LX, White PJ. 2013b. A conceptual model of root hair ideotypes for future agricultural environments: what combination of traits should be targeted to cope with limited P availability? Annals of Botany 112: 317–330.

Brundrett MC. 2021. Auditing data resolves systemic errors in databases and confirms mycorrhizal trait consistency for most genera and families of flowering plants. Mycorrhiza: 671–683.

Brundrett MC, Tedersoo L. 2018. Evolutionary history of mycorrhizal symbioses and global host plant diversity. New Phytologist 220: 1108–1115.

Brundrett M, Tedersoo L. 2019. Misdiagnosis of mycorrhizas and inappropriate recycling of data can lead to false conclusions. New Phytologist 221: 18–24.

Carminati A, Passioura JB, Zarebanadkouki M, Ahmed MA, Ryan PR, Watt M, Delhaize E. 2017. Root hairs enable high transpiration rates in drying soils. New Phytologist 216: 771–781.

Cosme M, Fernández I, van der Heijden MGA, Pieterse CMJ. 2018. Non-mycorrhizal plants: The exceptions that prove the rule. Trends in Plant Science: 1–11.

Dallstream C, Soper FM. 2024. Integrating edaphic gradients and community assembly concepts into the multidimensional root trait space. New Phytologist 243: 509–512.

Delhaize E, James RA, Ryan PR. 2012. Aluminium tolerance of root hairs underlies genotypic differences in rhizosheath size of wheat (Triticum aestivum) grown on acid soil. New Phytologist 195: 609–619.

Díaz S, Kattge J, Cornelissen JHC, Wright IJ, Lavorel S, Dray S, Reu B, Kleyer M, Wirth C, Prentice IC, et al. 2016. The global spectrum of plant form and function. Nature 529: 167–171.

Ding JX, Kong D, Zhang Z, Cai Q, Xiao J, Liu Q, Yin H. 2020. Climate and soil nutrients differentially drive multidimensional fine root traits in ectomycorrhizal-dominated alpine coniferous forests. Journal of Ecology 108: 2544–2556.

Dowle M, Srinivasan A. 2020. data.table: Extension of ‘data.fram’. https://cran.r-project.org/package=data.table.

Durka W, Michalski SG. 2012. Daphne: a dated phylogeny of a large European flora for phylogenetically informed ecological analyses. Ecology 93: 2297.

Eissenstat DM. 1992. Costs and benefits of constructing roots of small diameter. Journal of Plant Nutrition 15: 763–782.

Farquhar L. 1996. Root hairs: specialized tubular cells extending root surfaces. The Botanical Review 62: 1–40.

Fischer M, Bossdorf O, Gockel S, Hansel F, Hemp A, Hessenmoller D, Korte G, Nieschulze J, Pfeiffer S, Prati D, et al. 2010. Implementing large-scale and long-term functional biodiversity research: The Biodiversity Exploratories. Basic and Applied Ecology 11: 473–485.

Fort F, Cruz P, Catrice O, Delbrut A, Luzarreta M, Stroia C, Jouany C. 2015. Root functional trait syndromes and plasticity drive the ability of grassland Fabaceae to tolerate water and phosphorus shortage. Environmental and Experimental Botany 110: 62–72.

Freschet GT, Bellingham PJ, Lyver PO, Bonner KI, Wardle D a. 2013a. Plasticity in above- and belowground resource acquisition traits in response to single and multiple environmental factors in three tree species. Ecology and Evolution 3: 1065–1078.

Freschet GT, Cornelissen JHC, van Logtestijn RSP, Aerts R. 2010. Evidence of the ‘plant economics spectrum’ in a subarctic flora. Journal of Ecology 98: 362–373.

Freschet GT, Cornwell WK, Wardle DA, Elumeeva TG, Liu W, Jackson BG, Onipchenko VG, Soudzilovskaia NA, Tao J, Cornelissen JHC. 2013b. Linking litter decomposition of above- and below-ground organs to plant-soil feedbacks worldwide. Journal of Ecology 101: 943–952.

Freschet GT, Pagès L, Iversen C, Comas L, Rewald B, Roumet C, Klimešová J, Zadworny M, Poorter H, Postma J. 2021a. A starting guide to root ecology: Strengthening ecological concepts and standardizing root classification, sampling, processing and trait measurements. New Phytologist 232: 973–1122.

Freschet GT, Roumet C, Comas LH, Weemstra M, Bengough AG, Rewald B, Bardgett RD, De Deyn GB, Johnson D, Klimešová J, et al. 2021b. Root traits as drivers of plant and ecosystem functioning: current understanding, pitfalls and future research needs. New Phytologist 232: 1123–1158.

Gahoonia TS, Care D, Nielsen NE. 1997. Root hairs and phosphorus acquisition of wheat and barley cultivars. Plant and Soil 191: 181–188.

Gahoonia TS, Nielsen NE. 1998. Direct evidence on participation of root hairs in phosphorus (32P) uptake from soil. Plant and Soil 198: 147–152.

Garnier S. 2018. viridis: Default Color Maps from ‘matplotlib’. https://cran.r-project.org/package=viridis.

Gilroy S, Jones DL. 2000. Through form to function: Root hair development and nutrient uptake. Trends in Plant Science 5: 56–60.

Govindarajulu M, Pfeffer PE, Jin H, Abubaker J, Douds DD, Allen JW, Bücking H, Lammers PJ, Shachar-Hill Y. 2005. Nitrogen transfer in the arbuscular mycorrhizal symbiosis. Nature 435: 819–823.

Guerrero-Ramirez N et al. 2020. Global root traits (GRooT) database. Global Ecology and Biogeography 30: 25–37.

Guilbeault-Mayers X, Lambers H, Laliberté E. 2024. Coordination among leaf and fine-root traits along a strong natural soil fertility gradient. Plant and Soil.

Haling RE, Brown LK, Bengough AG, Young IM, Hallett PD, White PJ, George TS. 2013. Root hairs improve root penetration, root-soil contact, and phosphorus acquisition in soils of different strength. Journal of Experimental Botany 64: 3711–3721.

Haling RE, Yang Z, Shadwell N, Culvenor RA, Stefanski A, Ryan MH, Sandral GA, Kidd DR, Lambers H, Simpson RJ. 2016. Root morphological traits that determine phosphorus-acquisition efficiency and critical external phosphorus requirement in pasture species. Functional Plant Biology 43: 815–826.

Harrell FEJ. 2020. Hmisc: Harrell Miscellaneous. https://cran.r-project.org/package=Hmisc.

van der Heijden MGA, Martin FM, Selosse M-A, Sanders IR. 2015. Mycorrhizal ecology and evolution: the past, the present, and the future. New Phytologist 205: 1406–1423.

Hijmans RJ. 2020. raster: Geographic Data Analysis and Modeling. https://cran.r-project.org/package=raster.

Hill JO, Simpson RJ, Moore a. D, Chapman DF. 2006. Morphology and response of roots of pasture species to phosphorus and nitrogen nutrition. Plant and Soil 286: 7–19.

Hodge A, Fitter AH. 2010. Substantial nitrogen acquisition by arbuscular mycorrhizal fungi from organic material has implications for N cycling. Proceedings of the National Academy of Sciences 107: 13754–13759.

Hoeksema JD, Chaudhary VB, Gehring CA, Johnson NC, Karst J, Koide RT, Pringle A, Zabinski C, Bever JD, Moore JC, et al. 2010. A meta-analysis of context-dependency in plant response to inoculation with mycorrhizal fungi. Ecology Letters 13: 394–407.

Holdaway RJ, Richardson SJ, Dickie IA, Peltzer DA, Coomes DA. 2011. Species- and community-level patterns in fine root traits along a 120 000-year soil chronosequence in temperate rain forest. Journal of Ecology 99: 954–963.

Hope RM. 2013. Rmisc: Ryan Miscellaneous. https://cran.r-project.org/package=Rmisc.

Iversen CM, McCormack ML, Powell AS, Blackwood CB, Freschet GT, Kattge J, Roumet C, Stover DB, Soudzilovskaia NA, Valverde-Barrantes OJ, et al. 2017. A global Fine-Root Ecology Database to address below-ground challenges in plant ecology. New Phytologist 215: 15–26.

Jakobsen I, Chen B, Munkvold L, Lundsgaard T, Zhu Y. 2005. Contrasting phosphate acquisition of mycorrhizal fungi with that of root hairs using the root hairless barley mutant. : 928–938.

Kattge J, Bönisch G, Díaz S, Lavorel S, Prentice IC, Leadley P, Tautenhahn S, Werner GDA, Aakala T, Abedi M, et al. 2020. TRY plant trait database – enhanced coverage and open access. Global Change Biology 26: 119–188.

Kelley A. 1950. Mycotrophy in plants. Lectures on the biology of mycorrhizae and related structures. USA: Chronica Botánica Co., Waltham, Mass.

Kong D, Wang J, Zeng H, Liu M, Miao Y, Wu H, Kardol P. 2017. The nutrient absorption–transportation hypothesis: optimizing structural traits in absorptive roots. New Phytologist 213: 1569–1572.

Kumar A, Shahbaz M, Koirala M, Blagodatskaya E, Seidel SJ, Kuzyakov Y, Pausch J. 2019. Root trait plasticity and plant nutrient acquisition in phosphorus limited soil. Journal of Plant Nutrition and Soil Science 182: 945–952.

Lachaise T, Bergmann J, Rillig MC, van Kleunen M. 2021. Below- and aboveground traits explain local abundance, and regional, continental and global occurrence frequencies of grassland plants. Oikos 130: 110–120.

Lambers H, Raven JA, Shaver GR, Smith SE. 2008. Plant nutrient-acquisition strategies change with soil age. Trends in Ecology and Evolution 23: 95–103.

Lambers H, Teste FP. 2013. Interactions between arbuscular mycorrhizal and non-mycorrhizal plants: do non-mycorrhizal species at both extremes of nutrient availability play the same game? Plant, Cell and Environment Cell and Environment 36: 1911–1915.

Laughlin DC, Mommer L, Sabatini FM, Bruelheide H, Kuyper TW, McCormack ML, Bergmann J, Freschet GT, Guerrero-Ramírez NR, Iversen CM, et al. 2021. Root traits explain plant species distributions along climatic gradients yet challenge the nature of ecological trade-offs. Nature Ecology & Evolution 5: 1123–1134.

Ma X, Li X, Ludewig U. 2021. Arbuscular mycorrhizal colonization outcompetes root hairs in maize under low phosphorus availability. Annals of Botany 127: 155–166.

Ma X, Zarebanadkouki M, Kuzyakov Y, Blagodatskaya E, Pausch J, Razavi BS. 2018. Spatial patterns of enzyme activities in the rhizosphere: Effects of root hairs and root radius. Soil Biology and Biochemistry 118: 69–78.

Maherali H. 2014. Is there an association between root architecture and mycorrhizal growth response? New Phytologist 204: 192–200.

Maherali H. 2017. The evolutionary ecology of roots. New Phytologist 215: 1295–1297.

Martinez Arbizu P. 2019. pairwiseAdonis: Pairwise Multilevel Comparison using Adonis. R package version 0.3.

Marzec M, Melzer M, Szarejko I. 2015. Root hair development in the grasses: What we already know and what we still need to know. Plant Physiology 168: 407–414.

Matthus E, Zwetsloot M, Delory BM, Hennecke J, Andraczek K, Henning T, Mommer L, Weigelt A, Bergmann J. 2025. Revisiting the root economics space — its applications, extensions and nuances advance our understanding of fine-root functioning. Plant and Soil.

McCormack ML, Iversen CM. 2019. Physical and functional constraints on viable belowground acquisition strategies. Frontiers in Plant Science 10: 1215.

McGonigle TP, Miller MH, Evans DG, Fairchild GL, Swan JA. 1990. A new method which gives an objective-measure of colonization of roots. New Phytologist 115: 495–501.

Murdoch DJ, Chow ED. 2024. ellipse: Functions for drawing ellipses and ellipse-like confidence regions.

Nestler J, Wissuwa M. 2016. Superior root hair formation confers root efficiency in some, but not all, rice genotypes upon P deficiency. Frontiers in Plant Science 7: 1935.

Orme D, Freckleton R, Thomas D, Petzoldt T, Fritz S, Isaac N, Pearse W. 2018. caper: Comparative Analyses of Phylogenetics and Evolution in R. https://cran.r-project.org/package=caper.

Ostonen I, Püttsepp Ü, Biel C, Alberton O, Bakker MR, Lõhmus K, Majdi H, Metcalfe D, Olsthoorn AFM, Pronk A, et al. 2007. Specific root length as an indicator of environmental change. Plant Biosystems 141: 426–442.

Paradis E, Claude J, Strimmer K. 2004. APE: Analyses of phylogenetics and evolution in R language. Bioinformatics 20: 289–290.

Parasquive V, Brisson J, Guilbeault-Mayers X, Laliberté E, Chagnon PL. 2023. Contrasted root trait responses between saplings of an arbuscular and an ectomycorrhizal tree species in open field compared to forest conditions. Journal of Ecology 111: 1700–1710.

Price NS, Roncadori RW, Hussey RS. 1989. Cotton root growth as influenced by phosphorus nutrition and vesicular–arbuscular mycorrhizas. New Phytologist 111: 61–66.

R Core Team. 2020. R: A language and environment for statistical computing.

Reich PB. 2014. The world-wide ‘fast-slow’ plant economics spectrum: A traits manifesto. Journal of Ecology 102: 275–301.

Revell L. 2012. phytools: An R package for phylogenetic comparative biology (and other things). Methods in Ecology and Evolution 3: 217–223.

Rose L. 2017. Pitfalls in root trait calculations: How ignoring diameter heterogeneity can lead to overestimation of functional traits. Frontiers in Plant Science 8: 898.

Schweiger PF, Robson AD, Barrow NJ. 1995. Root hair length determines beneficial effect of a Glomus species on shoot growth of some pasture species. New Phytologist 131: 247–254.

Siqueira JO, Saggin-Júnior OJ. 2001. Dependency on arbuscular mycorrhizal fungi and responsiveness of some Brazilian native woody species. Mycorrhiza 11: 245–255.

Smith SE, Read DJ. 2008. Mycorrhizal Symbiosis. London: Academic Press.

Soetaert K. 2014. shape: Functions for plotting graphical shapes, colors. R package version 1.4.1.: http://CRAN.R-project.org/package=shape.

Soudzilovskaia NA, Vaessen S, Barcelo M, He J, Rahimlou S, Abarenkov K, Brundrett MC, Gomes SIF, Merckx V, Tedersoo L. 2020. FungalRoot: global online database of plant mycorrhizal associations. New Phytologist 227: 955–966.

Sun XG, Tang M. 2013. Effect of arbuscular mycorrhizal fungi inoculation on root traits and root volatile organic compound emissions of Sorghum bicolor. South African Journal of Botany 88: 373–379.

Warnes GR, Bolker B, Lumley T. 2020. gtools: Various R Programming Tools. https://cran.r-project.org/package=gtools.

Weigelt A, Mommer L, Andraczek K, Iversen CM, Bergmann J, Bruelheide H, Fan Y, Freschet GT, Guerrero-Ramírez NR, Kattge J, et al. 2021. An integrated framework of plant form and function: the belowground perspective. New Phytologist 232: 42–59.

Wen Z, Li H, Shen Q, Tang X, Xiong C, Li H, Pang J, Ryan MH, Lambers H, Shen J. 2019. Trade-offs among root morphology, exudation and mycorrhizal symbioses for phosphorus-acquisition strategies of 16 crop species.

Wen Z, White PJ, Shen J, Lambers H. 2022. Linking root exudation to belowground economic traits for resource acquisition. New Phytologist 233: 1620–1635.

Wickham H. 2010. ggplot2: elegant graphics for data analysis. Journal of Statistical Software 35.

Wickham H, François R, Henry L, Müller K. 2020a. dplyr: A grammar of data manipulation. R package version 1.0.2. https://cran.r-project.org/package=dplyr.

Wickham H, Hester J, Chang W. 2020b. devtools: Tools to make developing R packages easier. https://cran.r-project.org/package=devtools.

Wilke CO. 2024. cowplot: Streamlined plot theme and plot annotations for “ggplot2”.

Wright IJ, Reich PB, Westoby M, Ackerly DD, Baruch Z, Bongers F, Cavender-Bares J, Chapin T, Cornelissen JHC, Diemer M, et al. 2004. The worldwide leaf economics spectrum. Nature 428: 821–827.

Wu QS, Liu CY, Zhang DJ, Zou YN, He XH, Wu QH. 2016. Mycorrhiza alters the profile of root hairs in trifoliate orange. Mycorrhiza 26: 237–247.

Yang Z, Culvenor RA, Haling RE, Stefanski A, Ryan MH, Sandral GA, Kidd DR, Lambers H, Simpson RJ. 2015. Variation in root traits associated with nutrient foraging among temperate pasture legumes and grasses. Grass and Forage Science 72: 93–103.

Yu R, Li X, Xiao Z, Lambers H. 2020. Phosphorus facilitation and covariation of root traits in steppe species. New Phytologist 226: 1285–1298.

Zhao J, Guo B, Hou Y, Yang Q, Feng Z, Zhao Y, Yang X, Fan G, Kong D. 2024. Multi-dimensionality in plant root traits: progress and challenges. Journal of Plant Ecology 17.

Zhu J, Zhang C, Lynch JP. 2010. The utility of phenotypic plasticity of root hair length for phosphorus acquisition. Functional Plant Biology 37: 313–322.

