## Supporting Information for "Root hairs and mycorrhiza represent alternative phylogenetically conserved strategies for belowground absorptive surface maximization"

The following Supporting Information is available for this article:

**Fig. S1** Microscopic image of first order root of *Helictrotrichon pubescens* (Huds.) Domort., Poaceae, obligate mycorrhizal.

**Fig. S2** Pairwise correlations of all measured traits.

**Fig. S3** Extended phylogenetically informed principal component analysis.

**Fig. S4** Variation in mycorrhizal colonization (%M) between plants of different mycorrhizal status.

**Table S1** Phylogenetic signal of all measured traits.

**Table S2** Phylogenetically informed principal component analysis of root hair traits and %M.

**Table S3** Permanova based on pairwise dissimilarities of plant functional types and mycorrhizal types within the principal component analysis displayed in Table S2.

**Table S4** Extended phylogenetically informed principal component analysis.

**Table S5** Permanova based on pairwise dissimilarities of plant functional types and mycorrhizal types within the extended principal component analysis displayed in Table S4.

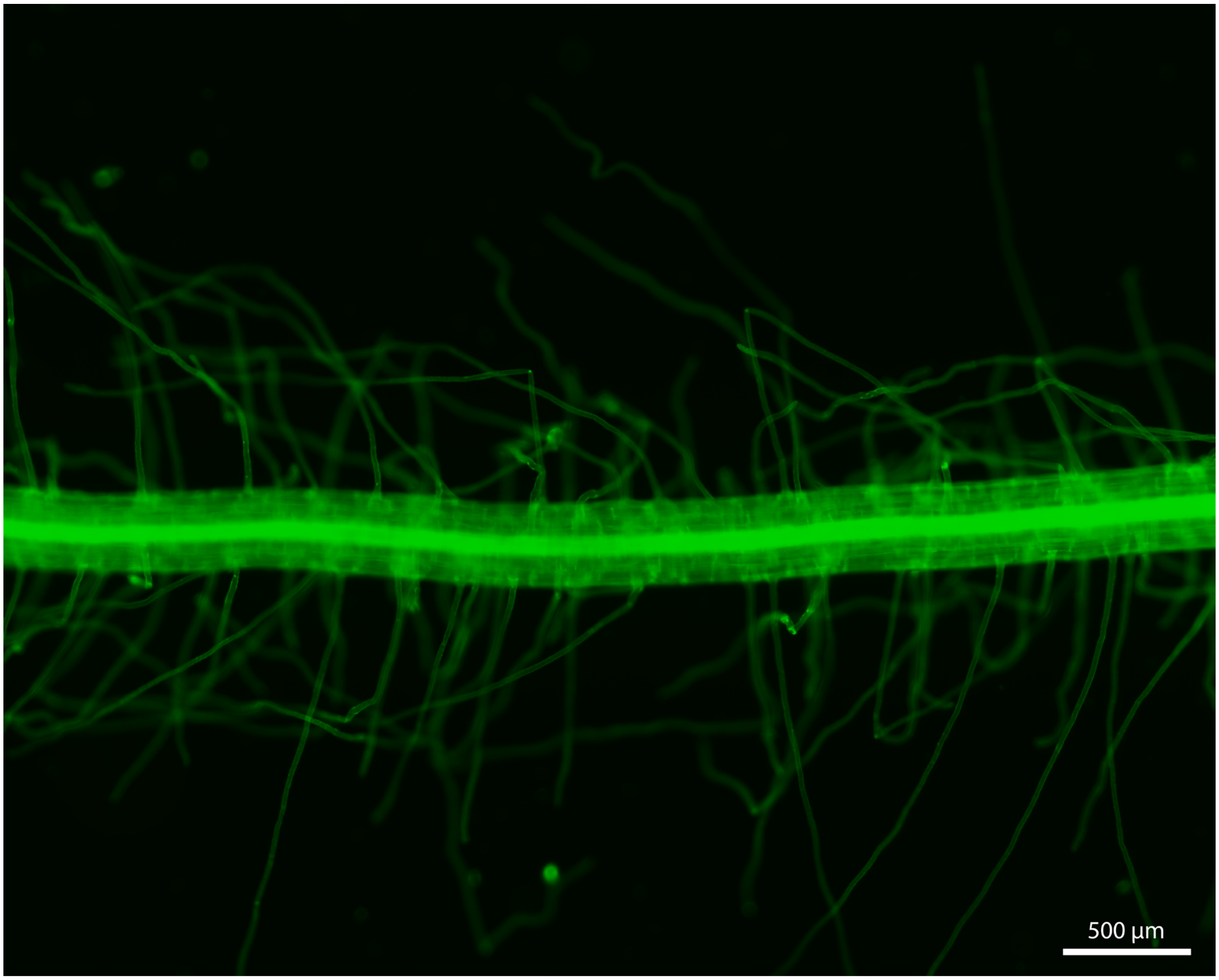


**Fig. S1 Microscopic image of first order root of *Helictrotrichon pubescens* (Huds.) Domort., Poaceae, obligate mycorrhizal.** Tissue outside of stele and stele itself are clearly distinguishable. The image was taken using a Zeiss AxioCam at a magnification of x50 using a 430 nm fluorescence filter.

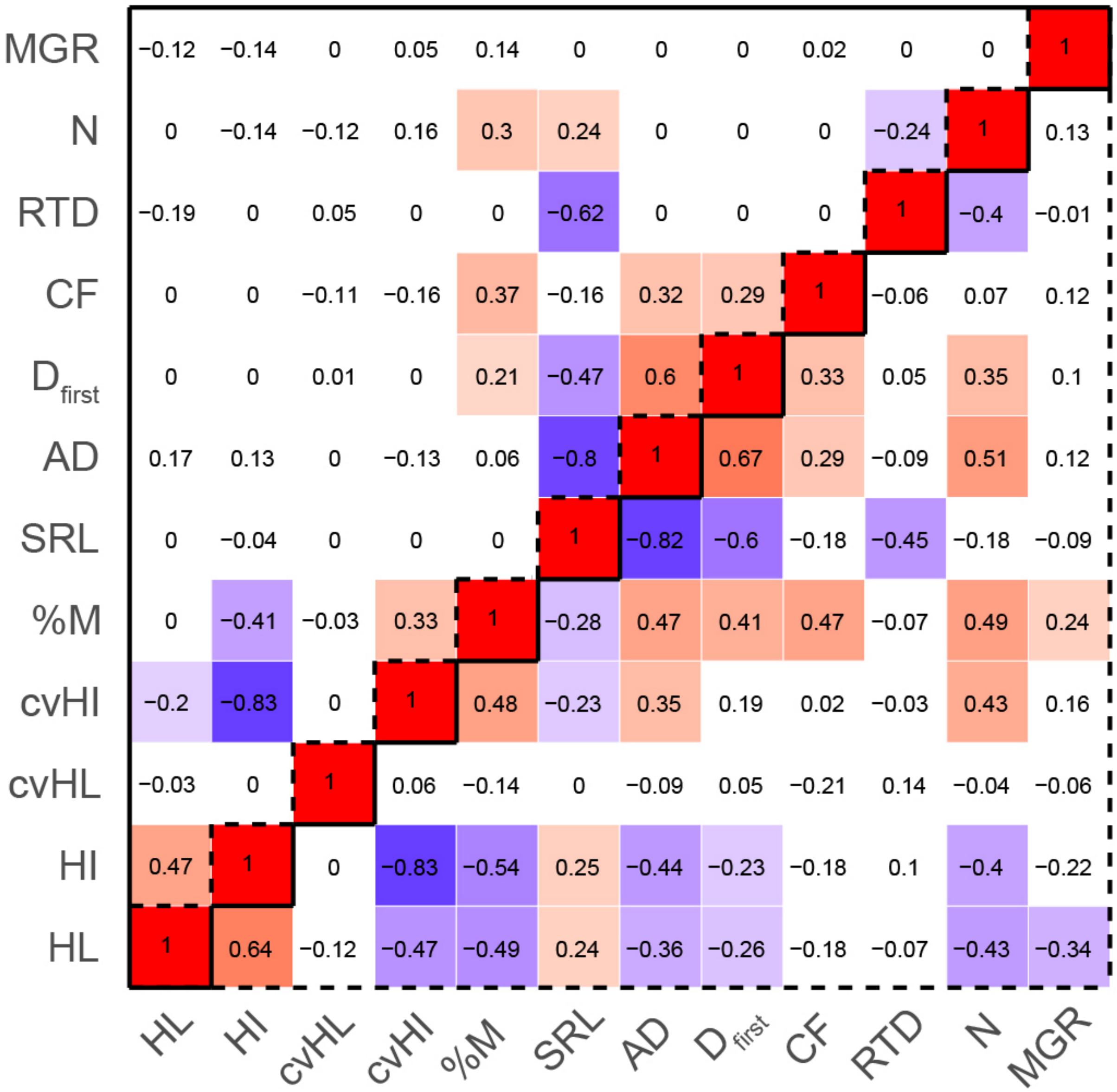


**Fig. S2 Pairwise correlations of all measured traits.** Upper triangle (solid line) represents phylogenetically informed correlations from PGLS (phylogenetic generalized least square) models while lower triangle (dashed line) represents correlations of raw traits. Displayed is the strength of the Pearson correlation while significant negative values ($P < 0.05$) appear in shades of blue and significant positive values appear in shades of red. HL – hair length, HI – hair incidence, cvHL – coefficient of variation in hair length, cvHI – coefficient of variation in hair incidence, %M - % mycorrhizal colonization, SRL – specific root length, AD – average diameter, $D_{first}$ – diameter of first order roots, CF – cortex fraction, RTD – root tissue density, N – root nitrogen content, MGR – mycorrhizal growth response.

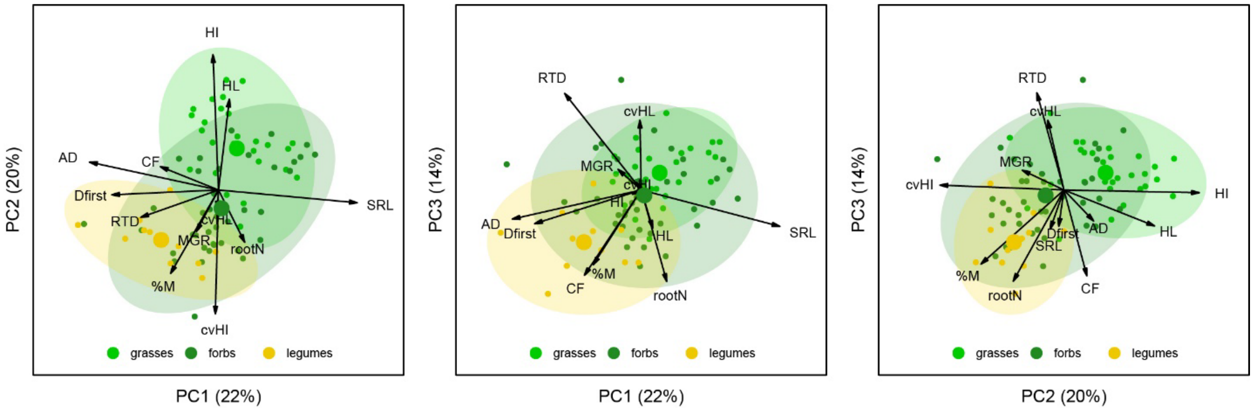


**Fig. S3 Extended phylogenetically informed principal component analysis.** Displayed are species based on their functional group (grasses, forbs, legumes). Ellipsoids and large dots display 95% confidence intervals and centroids. PCA results can be found in table S4. HL – hair length, HI – hair incidence, cvHL – coefficient of variation in hair length, cvHI – coefficient of variation in hair incidence, %M - % mycorrhizal colonization, SRL – specific root length, AD – average diameter, $D_{first}$ – diameter of first order roots, CF – cortex fraction, RTD – root tissue density, N – root nitrogen content, MGR – mycorrhizal growth response.

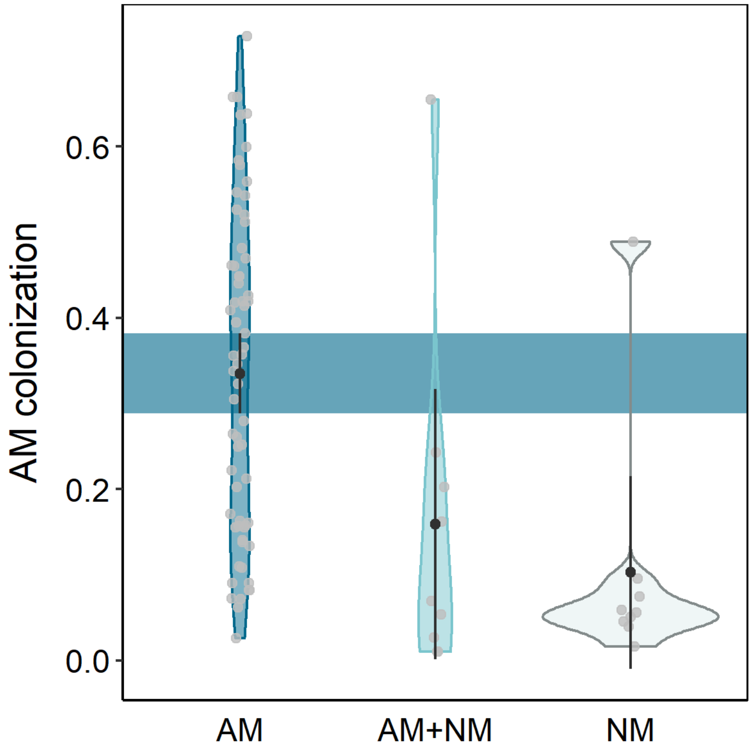


**Fig. S4 Variation in mycorrhizal colonization (%M) between plants of different mycorrhizal status.** Displayed are single species values with kernel density plots as well as mean (black dot) and 95% confidence interval per group. Non-overlapping confidence intervals are highlighted by a coloured ribbon to visualize group differences. AM – obligate mycorrhizal, AM+NM – facultative mycorrhizal, NM – non-mycorrhizal.

**Table S1 Phylogenetic signal of all measured traits.** Displayed is Pagel's lambda. HL – hair length, HI – hair incidence, cvHL – coefficient of variation in hair length, cvHI – coefficient of variation in hair incidence, %M - % mycorrhizal colonization, SRL – specific root length, AD – average diameter, $D_{first}$ – diameter of first order roots, CF – cortex fraction, RTD – root tissue density, N – root nitrogen content, MGR – mycorrhizal growth response.

| | **lambda** | ***P*** |
|---|---|---|
| **%M** | 0.831191 | 1.45E-12 |
| **HL** | 0.653663 | 2.78E-08 |
| **HI** | 0.789602 | 3.97E-18 |
| **SRL** | 0.733511 | 1.63E-05 |
| **AD** | 0.988914 | 2.86E-15 |
| **$D_{first}$** | 0.774926 | 1.95E-07 |
| **CF** | 0.656625 | 1.18E-05 |
| **RTD** | 0.220441 | 7.72E-02 |
| **N** | 0.995809 | 8.55E-22 |
| **cvHI** | 0.682502 | 3.36E-07 |
| **cvHL** | 0.768836 | 1.77E-02 |
| **MGR** | 0.687188 | 5.87E-04 |

**Table S2 Phylogenetically informed principal component analysis of root hair traits and %M**. lambda = 0.606. HL – hair length, HI – hair incidence, cvHL – coefficient of variation in hair length, cvHI – coefficient of variation in hair incidence, %M - percent mycorrhizal colonization.

| | **PC1** | **PC2** | **PC3** | **PC4** | **PC5** |
|---|---|---|---|---|---|
| **SD** | 1.496 | 1.065 | 0.871 | 0.812 | 0.456 |
| **Variance** | 0.448 | 0.227 | 0.152 | 0.132 | 0.042 |
| **HL** | 0.584 | -0.370 | 0.712 | 0.063 | 0.105 |
| **%M** | -0.591 | -0.475 | -0.052 | 0.649 | -0.028 |
| **HI** | 0.912 | 0.044 | -0.078 | 0.213 | -0.338 |
| **cvHI** | -0.835 | -0.008 | 0.407 | -0.238 | -0.283 |
| **cvHL** | -0.128 | 0.878 | 0.280 | 0.365 | 0.044 |

**Table S3 Permanova based on pairwise dissimilarities of plant functional types and mycorrhizal types within the principal component analysis displayed in Table S3.** AM – obligate mycorrhizal, AM+NM – facultative mycorrhizal, NM – non-mycorrhizal.

| | | | **Sums of Squares** | ***F*** | ***R²*** | ***P*** |
|---|---|---|---|---|---|---|
| **AM+NM** | **vs** | **AM** | 1260.751 | 2.373 | 0.033 | 0.110 |
| **AM+NM** | **vs** | **NM** | 785.512 | 3.516 | 0.180 | 0.066 |
| **AM** | **vs** | **NM** | 4500.254 | 8.552 | 0.109 | 0.003 |
| **grasses** | **vs** | **forbs** | 10561.737 | 26.129 | 0.278 | 0.002 |
| **grasses** | **vs** | **legumes** | 9796.955 | 34.155 | 0.473 | 0.002 |
| **forbs** | **vs** | **legumes** | 887.421 | 2.139 | 0.041 | 0.137 |

**Table S4 Extended phylogenetically informed principal component analysis**, lambda = 0.588. HL – hair length, HI – hair incidence, cvHL – coefficient of variation in hair length, cvHI – coefficient of variation in hair incidence, %M - % mycorrhizal colonization, SRL – specific root length, AD – average diameter, $D_{first}$ – diameter of first order roots, CF – cortex fraction, RTD – root tissue density, N – root nitrogen concentration, MGR – mycorrhizal growth response.

| | **PC1** | **PC2** | **PC3** | **PC4** | **PC5** | **PC6** | **PC7** | **PC8** | **PC9** | **PC10** | **PC11** | **PC12** |
|---|---|---|---|---|---|---|---|---|---|---|---|---|
| **SD** | 1.614 | 1.550 | 1.316 | 1.079 | 0.988 | 0.863 | 0.835 | 0.778 | 0.675 | 0.639 | 0.437 | 0.134 |
| **Variance** | 0.217 | 0.200 | 0.144 | 0.097 | 0.081 | 0.062 | 0.058 | 0.050 | 0.038 | 0.034 | 0.016 | 0.001 |
| **SRL** | 0.918 | -0.085 | -0.241 | 0.020 | -0.191 | 0.049 | 0.085 | -0.008 | 0.191 | 0.010 | 0.012 | 0.095 |
| **AD** | -0.852 | 0.184 | -0.192 | -0.136 | 0.099 | 0.086 | -0.070 | 0.251 | -0.246 | 0.181 | 0.014 | 0.076 |
| **$D_{first}$** | -0.700 | -0.031 | -0.221 | -0.320 | -0.228 | 0.167 | -0.085 | 0.096 | 0.503 | -0.078 | -0.032 | -0.006 |
| **HL** | 0.071 | 0.592 | -0.235 | -0.194 | 0.421 | 0.446 | -0.002 | -0.397 | -0.023 | -0.070 | -0.111 | 0.006 |
| **CF** | -0.372 | 0.149 | -0.559 | 0.336 | -0.396 | -0.011 | 0.346 | -0.076 | -0.163 | -0.323 | -0.030 | 0.002 |
| **%M** | -0.313 | -0.546 | -0.487 | 0.069 | 0.056 | -0.167 | 0.180 | -0.413 | 0.072 | 0.343 | 0.073 | -0.005 |
| **HI** | -0.037 | 0.895 | -0.018 | 0.033 | -0.138 | -0.118 | -0.188 | -0.151 | 0.050 | -0.012 | 0.319 | 0.000 |
| **RTD** | -0.504 | -0.177 | 0.637 | 0.129 | 0.169 | -0.317 | -0.056 | -0.311 | 0.089 | -0.226 | -0.044 | 0.054 |
| **N** | 0.167 | -0.333 | -0.598 | -0.226 | -0.036 | -0.242 | -0.584 | -0.084 | -0.135 | -0.154 | -0.051 | -0.001 |
| **cvHI** | -0.021 | -0.818 | 0.032 | -0.217 | 0.213 | 0.324 | 0.084 | 0.021 | -0.080 | -0.233 | 0.253 | 0.004 |
| **cvHL** | -0.009 | -0.104 | 0.449 | -0.482 | -0.643 | 0.168 | -0.012 | -0.259 | -0.194 | 0.088 | -0.031 | 0.002 |
| **MGR** | -0.139 | -0.260 | 0.123 | 0.734 | -0.183 | 0.418 | -0.372 | -0.072 | 0.020 | 0.077 | -0.002 | 0.003 |

**Table S5 Permanova based on pairwise dissimilarities of plant functional types and mycorrhizal types within the extended principal component analysis displayed in Table S5.** AM – obligate mycorrhizal, AM+NM – facultative mycorrhizal, NM – non-mycorrhizal.

| | | | Sums of Squares | *F* | $R^2$ | *P* |
|---|---|---|---|---|---|---|
| **AM+NM** | **vs** | **AM** | 6638.494 | 5.447 | 0.073 | 0.005 |
| **AM+NM** | **vs** | **NM** | 1522.688 | 1.512 | 0.092 | 0.178 |
| **AM** | **vs** | **NM** | 7206.628 | 6.139 | 0.081 | 0.005 |
| **grasses** | **vs** | **forbs** | 13089.700 | 13.567 | 0.168 | 0.001 |
| **grasses** | **vs** | **legumes** | 26578.000 | 39.506 | 0.510 | 0.001 |
| **forbs** | **vs** | **legumes** | 10684.200 | 9.732 | 0.166 | 0.001 |
